# Relationship Between Therapeutic Activity and Preferential Targeting of Toxic Soluble Aggregates by Amyloid-Beta-Directed Antibodies

**DOI:** 10.1101/2024.04.20.590412

**Authors:** Johanne M. Kaplan, Ebrima Gibbs, Juliane Coutts, Beibei Zhao, Ian Mackenzie, Neil R. Cashman

**Author notes:** Corresponding authors: Johanne Kaplan & Neil Cashman ProMIS Neurosciences, One Broadway, Cambridge, MA 02142.

## Abstract

**Background:** Amyloid-beta (Aβ)-directed antibodies tested clinically for therapeutic activity against Alzheimer’s disease (AD) have shown varying degrees of efficacy. Although all of these antibodies target the Aβ peptide, their binding profile to different molecular species of Aβ differs and may underlie the observed variability in clinical outcomes.

**Objective:** Explore the relationship between targeting of soluble toxic Aβ species and therapeutic efficacy.

**Methods:** Surface plasmon resonance (SPR) was used to conduct a side-by-side comparison of the binding of various Aβ-directed antibodies to monomers and soluble Aβ oligomers from AD brains. Immunohistochemistry was performed to assess reactivity with plaque. Preclinical activity was assessed in human amyloid precursor protein (APP) transgenic mouse models of AD.

**Results:** Non-selective, pan-Aβ reactive antibodies such as crenezumab and gantenerumab, which have failed to produce a clinical benefit, bound all forms of Aβ tested. In a competition assay, these antibodies lost the ability to bind toxic AD brain oligomers when exposed to monomers. Aggregate-selective antibodies such as aducanumab, lecanemab and donanemab, showed reduced monomer binding and a greater ability to withstand monomer competition which correlated with their reported inhibition of cognitive decline. Of the antibodies in earlier stages of clinical testing, ACU193 and PMN310 displayed the greatest ability to retain binding to toxic AD brain oligomers while PRX h2731 was highly susceptible to monomer competition. Plaque binding was observed with all aggregate-reactive antibodies with the exception of PMN310, which was strictly selective for soluble oligomers. Targeting of oligomers by PMN310 protected cognition and was not associated with microhemorrhages in mouse models of AD.

**Conclusions:** Overall, these results suggest that selectivity for soluble toxic Aβ oligomers may be a driver of clinical efficacy, with a potentially reduced risk of ARIA if engagement with plaque is minimized.

## INTRODUCTION

Strong genetic and experimental evidence supports a causative role for amyloid-beta (Aβ) in the pathogenesis of Alzheimer’s disease (AD) in agreement with the demonstrated ability of Aβ-directed antibodies such as aducanumab, lecanemab and donanemab to slow cognitive decline in pivotal clinical trials [1–3]. What remains unclear is the exact nature of the Aβ species being targeted by various antibodies and their relative contribution to clinical efficacy. While there exists a seeming correlation between cognitive benefit and antibody-mediated plaque clearance as measured by PET imaging [4], the same antibodies can also mediate the clearance of soluble toxic Aβ species such as oligomers. Because there are currently no available standardized methods for clinical measurement of these toxic Aβ species, it has been difficult to evaluate potential correlations with the efficacy of treatment.

A large body of experimental and clinical evidence suggests that soluble toxic Aβ aggregates, rather than insoluble fibrils and plaque, are the primary drivers of synaptic dysfunction, neuronal loss and cognitive decline in AD patients [5–10]. For example, genetic data indicate that the APP Arctic E693G, APP Osaka (E693Δ), and APP p.D678N mutations cause rapidly progressive dementia with abundant soluble toxic Aβ species, and little or no Aβ plaque as measured by PiB PET [11–13]. Accordingly, the intended mechanism of action for antibodies such as lecanemab, ACU193 and PMN310 is targeting of these toxic soluble species. Since soluble aggregates of Aβ are heterogeneous, questions also arise as to which subset(s) of aggregates are being targeted and how these antibodies might achieve efficacy. In this study, a side-by-side comparison of the binding profile of various Aβ-targeted antibodies to monomers, low and high molecular weight AD brain oligomers, and plaque was conducted and compared with known clinical outcomes. The results suggest that clinical efficacy may be related to both the ability of Aβ-directed antibodies to bind soluble toxic Aβ aggregates and a degree of specificity allowing for binding soluble toxic Aβ aggregates in the presence of competing Aβ monomers.

## MATERIALS & METHODS

### Reagents

The monoclonal antibodies solanezumab, crenezumab, gantenerumab, aducanumab, lecanemab, donanemab, ACU193 and PRX h2731 were produced by Creative Biolabs (Upton, NY, USA) based on published sequences. PMN310, a humanized IgG1 monoclonal antibody selective for toxic Aβ oligomers [14] was manufactured by KBI Biopharma (Durham, NC, USA). A mouse IgG2a version of PMN310 (mPMN310) was produced by Wuxi Biologics (Hong Kong, China). Hexafluoroisopropanol (HFIP)-treated recombinant Aβ42 and Aβ40 peptides (rPeptide, Watkinsville, GA, USA) were reconstituted in dimethyl sulfoxide (DMSO) (Sigma-Aldrich Canada, Oakville, ON) to give a stock concentration of 5 mM. Oligomers of Aβ42 were prepared by diluting the peptide solution in phenol red-free F12 medium (Thermo Fisher, Waltham MA, USA) to a final concentration of 100 µM and incubated for 24 h at 4°C followed by immediate use or storage at −80°C. Aβ40, which is less prone to aggregation, was used in monomer competition assays. Human IgG1 isotype control was purchased from BioLegend (San Diego, CA, USA).

### Brain extracts

Brain tissues from 15 different human AD patients were provided by Dr. Ian Mackenzie at the University of British Columbia, Vancouver, Canada. Informed consent for tissue collection at autopsy and neurodegenerative research use was obtained from the legal representative in accordance with local institutional review boards. Donor characteristics are summarized in supplemental Table 1. The diagnosis of AD was based on NINCDS-ADRDA clinical and pathological criteria. Samples from frontal cortex were weighed and submersed in ice-cold Tris-buffered saline (TBS) (20% w/v) with EDTA-free protease inhibitor cocktail (Roche Diagnostics, Laval QC, Canada), and homogenized using an Omni tissue homogenizer (Omni International Inc, Keenesaw GA, USA), 3 x 30 sec pulses with 30 sec pauses in between, all performed on ice. Homogenates were then subjected to ultracentrifugation at 70,000xg for 90 min. Supernatants (soluble extracts) were collected, aliquoted and stored at −80°C. The protein concentration was determined using a bicinchoninic acid (BCA) protein assay. Pools of brain extracts from 3-5 patients were used in each analysis.

### Size exclusion chromatography

Pooled soluble brain extracts were injected at 0.5 ml/min through a Superdex 75 (10/300) HPLC column (GE Healthcare Life Sciences, Pittsburg PA, USA) for 50 min and 0.25 ml fractions were collected. Molecular weight (MW) markers (Bio-Rad Laboratories, Mississauga ON, Canada) were run separately. Protein peaks were monitored by absorbance at O.D. 280 nm. Hydrodynamic fractions corresponding to a MW of ∼8kDa to ∼70kDa were pooled into a low molecular weight (LMW) fraction. Aβ monomers (MW ∼4.5kDa) were excluded from the LMW fraction. Fractions corresponding to a MW of >140kDa to ∼700kDa were pooled into a high molecular weight (HMW) fraction. The LMW and HMW fractions were concentrated, and total protein concentration was determined in a BCA assay. The fractions were then diluted to 100 μg/ml in phosphate-buffered saline, 3 mM EDTA, 0.05% surfactant P20 (PBS-EP) (GE Healthcare Life Sciences, Pittsburg PA, USA) containing BSA (2 mg/ml) for surface plasmon resonance (SPR) analysis.

### Surface plasmon resonance analysis

Surface plasmon resonance measurements were performed using a Molecular Affinity Screening System (MASS-2) instrument (Bruker Daltronics, Billerica, MA, USA). Antibodies were immobilized on high amine capacity sensor chips at a density of approximately 9,000-11,000 response units (RUs). To assess the binding of antibodies to native AβO, the LMW and HMW size-exclusion chromatography fractions of pooled soluble human AD brain extracts (100 µg/ml) were injected over the immobilized antibodies at 10 µl/min for 8 min. The binding responses from the resultant sensorgrams were double-referenced against unmodified reference surfaces and blank buffer injections. Binding response units (BRUs) were collected 30 sec post-injection stop during the dissociation period. In monomer competition studies, Aβ40 monomers (0.08 μM - 5 μM) were pre-injected over immobilized antibodies at 12.5 μl/min for 15 min followed by injection of brain extract (100 μg/ml) at 12.5 μl/min for 10 min. Percent binding response was calculated as: [(BRU) with monomers) / (BRU without monomers)] X100.

### Histology

Fresh-frozen, frontal cortex AD brain sections (25 μm) were mounted onto charged glass slides and fixed by immersion in −20 °C acetone for 20 min. Between subsequent steps, slides were washed extensively in Tris-buffered saline (TBS, pH 7.4) containing 0.2% Triton-X-100. To suppress endogenous peroxidase activity, sections were treated with 0.3% H_2_0_2_, followed by incubation in TBS buffer containing 10% normal goat serum (NGS), 3% BSA, and 0.2% Triton-X-100 to block non-specific binding sites. Next, an avidin/biotin blocking kit (Vector Laboratories, Burlington, ON, Canada) was employed as per the manufacturer’s instructions. Endogenous immunoglobulins were blocked by incubating tissue in Fab fragment goat anti-human IgG (H+L) (Jackson ImmunoResearch Labs, West Grove, PA, USA; 100 μg/ml) for 2 h at room temperature. Primary antibodies (solanezumab, gantenerumab, aducanumab, lecanemab, donanemab, ACU193, PRX h2731 and PMN310), were adjusted to 1 μg/ml in background-reducing antibody diluent (Agilent, Santa Clara, CA, USA) and applied overnight at 4 °C in a humidified chamber. Biotin-conjugated goat anti-human IgG (Fab’)2 antibody (Abcam, Waltham, MA, USA; 1 µg/ml) was subsequently added for 1 h at room temperature. Staining was visualized using the ABC method, with a Vectastain kit (Vector Laboratories) and diaminobenzidine (DAB) as the chromogen. Nuclei were counterstained using methyl green or hematoxylin QS (Vector Laboratories) and sections were dehydrated, cleared, and coverslipped.

Brain sections from knock-in *APP^SAA^*mice (*B6.Cg-App^tm1.1Dnli^/J* strain; Jackson Labs, Bar Harbor, ME), a mouse model of AD, were cut at a nominal thickness of 4 µm to 5 µm and stained with Perls’ Prussian Blue stain to detect the presence of hemosiderin as an indicator of hemorrhage in the brain. An unrelated stock spleen sample was used as a positive control to validate the staining procedure (supplemental Figure 1). The mice were part of a toxicology study conducted at Sequani (Herefordshire, UK) in compliance with good laboratory practice (GLP) and in accordance with the OECD principles of GLP. Animals were dosed once weekly for 26 weeks by subcutaneous injection with vehicle or 800 mg/kg of mPMN310 into alternate four-hand flank regions (upper/lower, left/right). Stained sections from 29 vehicle control mice and 29 mPMN310-treated mice were examined (14 males and 15 females in each group). The presence of Aβ plaque in these mice was confirmed by Amylo-Glo staining according to manufacturer’s instructions (Biosensis, Temecula, CA, USA)

### Cognition assessment in AD mouse model

Testing of mPMN310 in mice transgenic (Tg) for a single clinical mutant of the human amyloid precursor protein (hAPP [V717I]), expressed under control of the murine Thy1.2 promoter for neuron-specific expression, was conducted at reMYND (Leuven, Belgium) in compliance with standards for animal care and use. Starting at 5 months of age, hAPP-Tg mice received weekly intraperitoneal injections of mPMN310 (30 mg/kg) or vehicle for 6 weeks (N=17/group). Age-matched, non-Tg, wild-type mice injected with vehicle were included as a control group (N=17). At the end of the dosing period, cognition was assessed in a Morris Water Maze test in which mice placed in a pool of water were trained to locate a hidden platform in 4 training sessions of 4 trials over 4 consecutive days. The time (escape latency) and distance (search path) each mouse needed to locate the platform were measured. Results are shown as the mean ± standard error of the mean (SEM) of the 4 trials for each training session.

### Statistical analysis

Statistical analysis was performed with GraphPad Prism 7 as described in the figure legends.

## RESULTS

### Aβ-directed antibodies bind to soluble species in AD brain extract

The neurotoxicity of soluble Aβ aggregates of various sizes ranging from dimers to protofibrils has been described *in vitro* and *in vivo* by multiple investigators [7, 15–17]. Preferential targeting of these soluble species, as opposed to insoluble fibrils, has been proposed as the primary mechanism underlying lecanemab’s clinical efficacy [3, 18]. We therefore compared the binding of lecanemab and other clinical-stage antibodies to soluble species of Aβ derived from AD brains using surface plasmon resonance (SPR). Equivalent amounts of antibodies were immobilized on a sensor chip and soluble AD brain extract fractionated by size-exclusion chromatography into a low molecular (LMW) fraction excluding monomers (∼8-70 kDa, oligomer-enriched), and a high molecular weight (HMW) fraction (∼140-700 kDa, protofibril-enriched), were injected over the surface (Fig. 1). As shown in Figure 2, lecanemab displayed binding to both soluble AD brain aggregate fractions in agreement with previous reports showing strong binding of lecanemab to oligomers (defined by the authors as <75 kDa), small protofibrils (defined as 75-400 kDa) and large protofibrils (defined as 300-5,000 kDa) [19, 20]. The same reports indicated that the Aβ species with the most toxicity resided in the 20-80 kDa and 80-500 kDa fractions, overlapping with the LMW and HMW fractions tested herein thereby suggesting that reactivity against both fractions may contribute to the protective activity of lecanemab. While emphasis has been placed on the reactivity of lecanemab with protofibrils over 75 kDa, the binding profile of this antibody to Aβ species in AD brain appears broader and it likely targets a range of toxic soluble species (particularly LMW oligomers) of various sizes as opposed to a specific subpopulation with a defined molecular weight cut-off.

**Figure 1.**
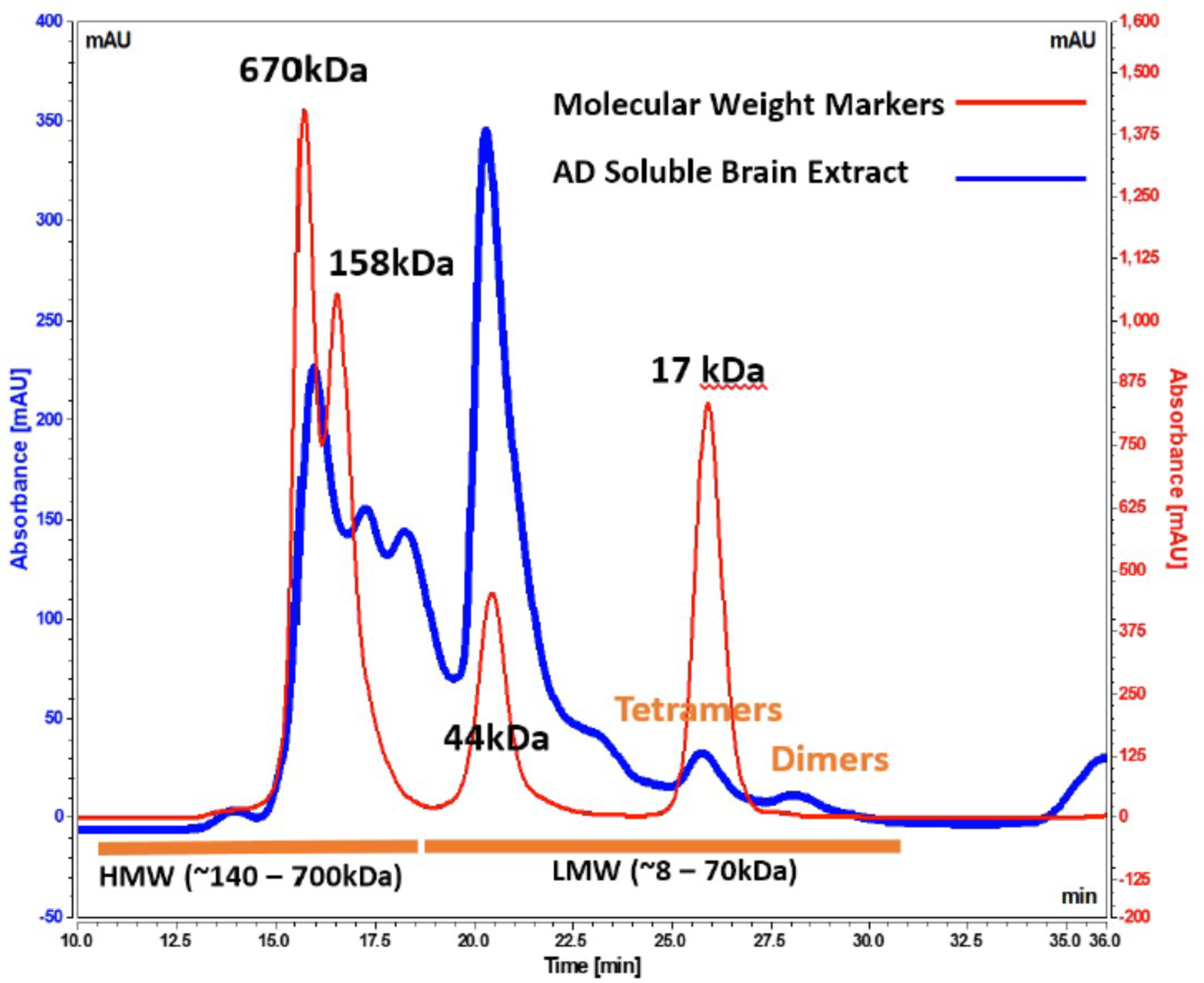
Size-exclusion chromatogram of soluble AD brain extract. Fractionated soluble brain extract was pooled into a LMW fraction of ∼8-70 kDa and a HMW fraction of ∼140-700 kDa. Aβ monomers (4.5 kDa) were excluded.

**Figure 2.**
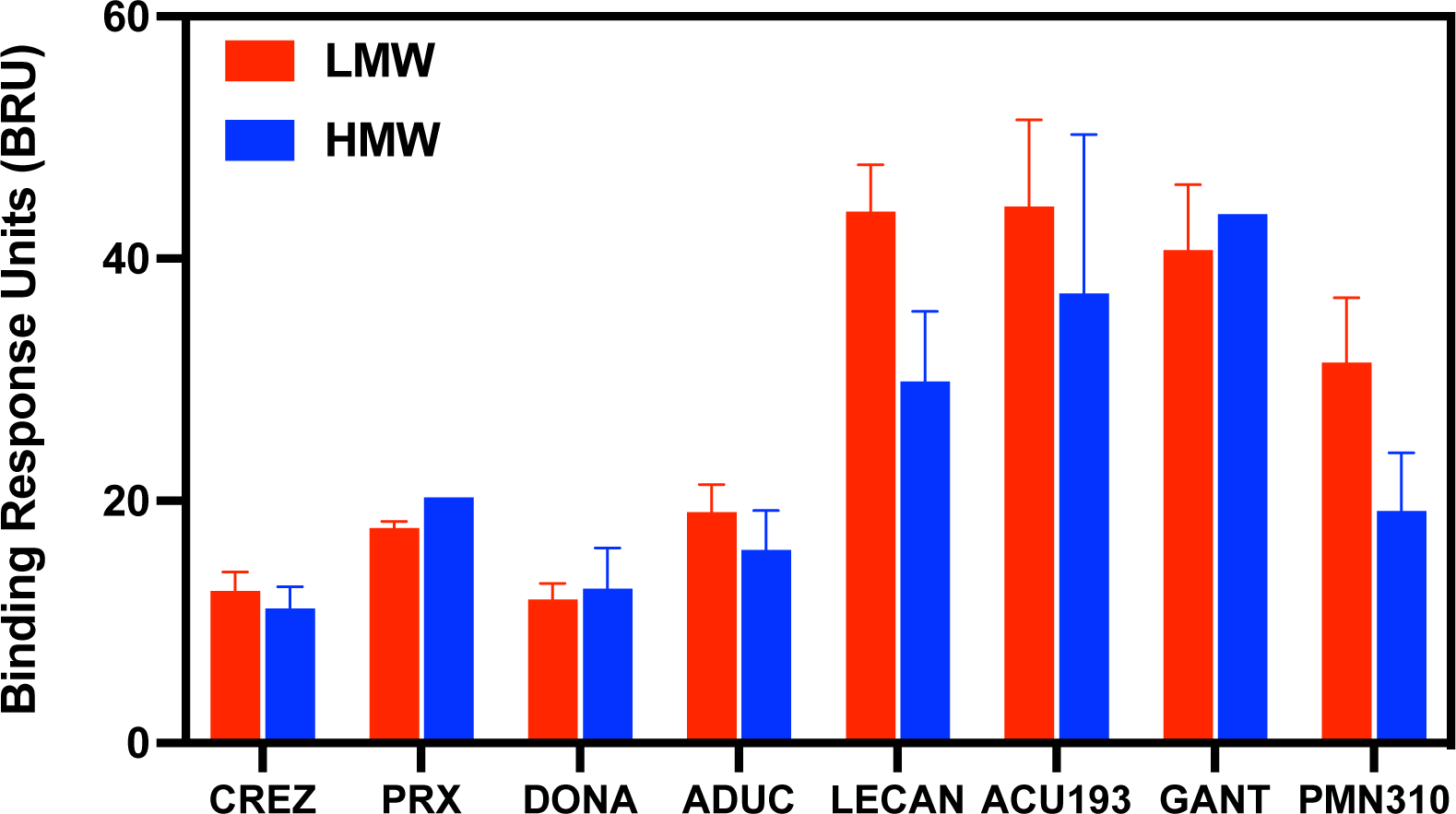
Binding of Aβ-directed antibodies to soluble species in AD brain extract. SPR measurement of immobilized antibody binding to the LMW and HMW fractions of soluble AD brain extract. Results shown are the mean ± SEM of combined data from 11 independent studies for the LMW fraction and 7 studies for the HMW fraction. Not all antibodies were tested in all studies. CREZ: crenezumab (LMW - 9 studies, HMW - 5 studies), PRX: PRX h2731 (LMW - 2 studies, HMW - 1 study), DONA: donanemab (LWW - 11 studies, HMW - 7 studies), ADUC: aducanumab (LMW - 11 studies, HMW - 7 studies), LECAN: lecanemab (LMW - 11 studies, HMW - 6 studies), ACU193 (LMW - 10 studies, HMW - 6 studies), GANT: gantenerumab (LMW - 5 studies, HMW - 1 study), PMN310 (LMW - 11 studies, HMW - 7 studies). Multiple paired *t*-tests for LMW *vs* HMW, *p* = 0.05 and 0.09 for LECAN and PMN310, respectively. *p* > 0.3 for all other antibodies.

Other antibodies (crenezumab, PRX h2731, donanemab, aducanumab and ACU193) also showed comparable levels of binding to soluble species in both the LMW and HMW fractions of AD brain extract (Fig. 2). PMN310, like lecanemab, showed binding to both fractions with a trend for preferential reactivity with the LMW toxic oligomer-enriched fraction.

### Ability to avoid monomer competition corresponds with clinical efficacy

Clinical results obtained to date indicate that therapeutic efficacy is significantly affected by the binding profile of Aβ-directed antibodies [21]. Antibodies with pan-reactivity such as gantenerumab and crenezumab which bind all species of Aβ, including toxic oligomers, did not achieve clinical benefit [22, 23]. These results suggest that efficacy may depend on antibody selectivity and the ability to avoid interaction with abundant, non-toxic monomers, which can reduce the effective dose available for targeting pathogenic soluble species present at concentrations at least 1,000 fold lower than monomers in the brain [16, 24–26]. This concept was explored in SPR binding studies to determine whether a relationship exists between positive clinical data and the ability of an antibody to retain binding to soluble oligomers from AD brain extract in the face of monomer competition.

First, the binding of Aβ-directed antibodies to increasing concentrations of Aβ monomers (0.08-5.0 μM) was measured by SPR (Fig. 3). Pan-Aβ reactive antibodies such as crenezumab, gantenerumab and PRX h2731 showed high levels of binding, even at the lowest concentration of monomers tested. Antibodies with more selectivity for aggregated forms of Aβ such as donanemab, aducanumab and lecanemab showed markedly lower levels of monomer binding, although lecanemab still displayed relatively robust binding at the highest concentrations of monomers. PMN310 and ACU193, both designed to selectively target oligomers, showed low monomer binding across the range of concentrations.

**Figure 3.**
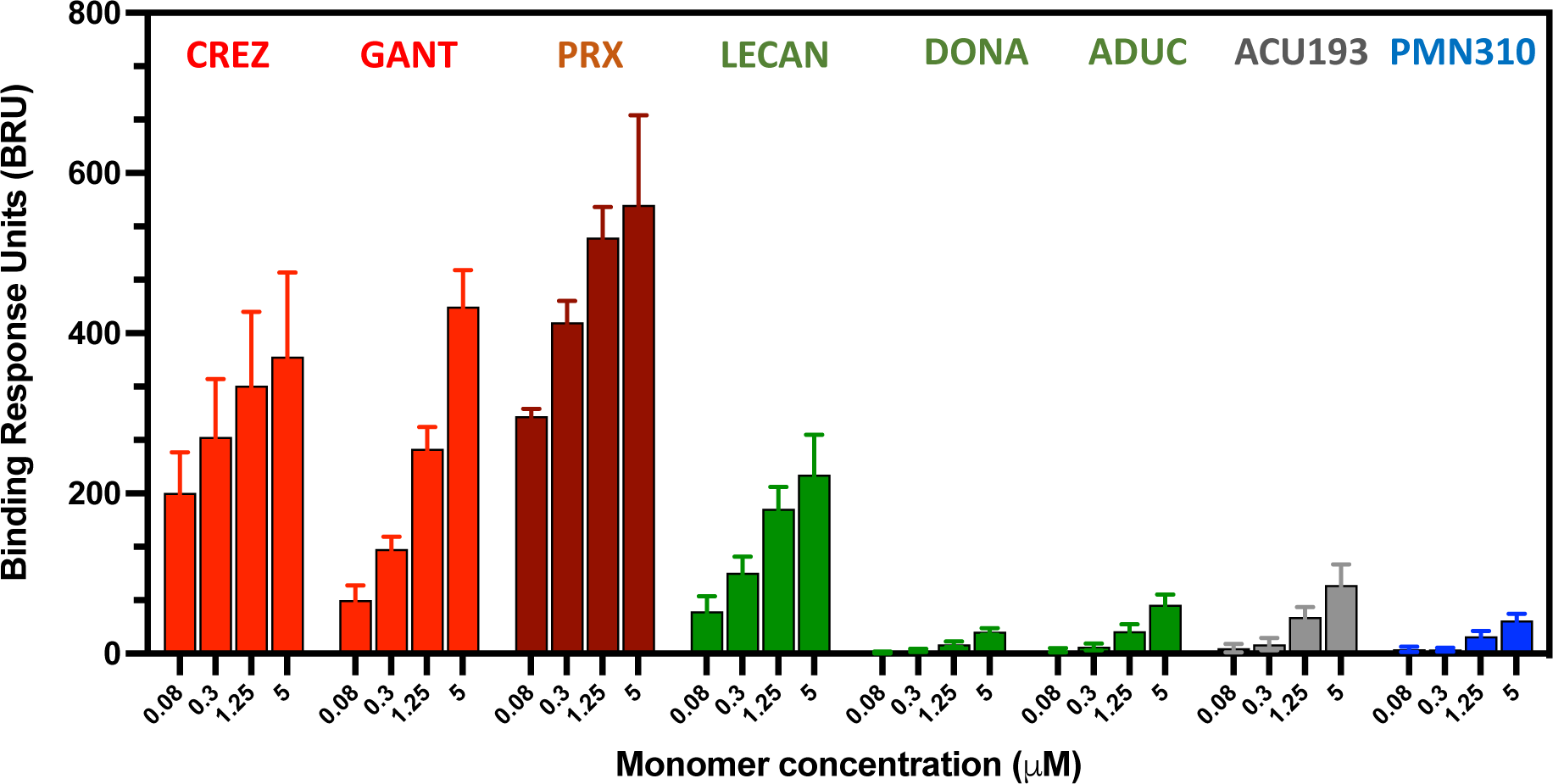
Binding of Aβ-directed antibodies to monomers. SPR measurement of immobilized antibody binding to Aβ monomers. Results shown are the mean ± SEM of combined data from 6 independent studies. Not all antibodies were tested in all studies. CREZ and GANT - 4 studies, PRX - 2 studies, LECAN, DONA, ADUC, ACU193 and PMN310 - 6 studies.

Binding of antibodies to the LMW and HMW fractions from soluble AD brain extract was then measured either in the absence of Aβ monomers (100% binding response) or in the presence of increasing concentrations of monomers to mimic the competition that antibodies will encounter in the circulation and in the brain after systemic administration. Binding to the LMW fraction of soluble AD brain extract is of particular interest since it has been reported to be enriched for toxic oligomers as assessed by inhibition of long-term potentiation [9], induction of cognitive deficit in rats [6, 27], and activation of microglia [28]. In side-by-side comparison studies, binding of oligomers by antibodies with pan-Aβ reactivity such as crenezumab, gantenerumab and PRX h2731 was strongly inhibited by low concentrations of monomers thereby reducing the amount of antibody available to bind oligomers in the LMW (Fig. 4a) or HMW (Fig. 4b) fractions of soluble AD brain extract. In contrast, antibodies with greater selectivity for aggregated Aβ were better able to withstand monomer competition and retained measurable binding to LMW and HMW oligomers even at high concentrations of monomers.

**Figure 4.**
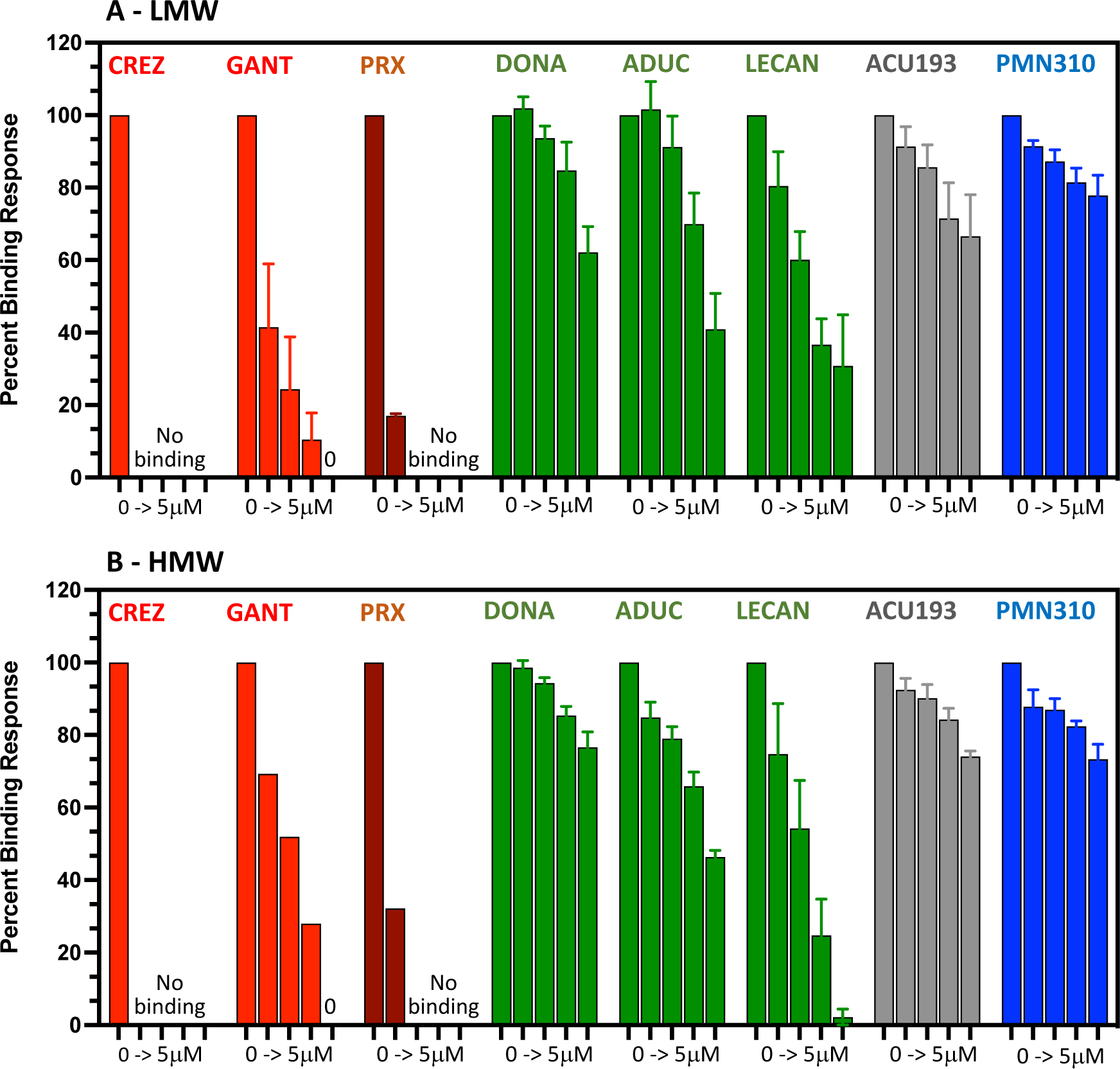
Binding to LMW and HMW brain oligomers in the face of monomer competition. SPR measurement of immobilized antibody binding to the LMW (A) or HMW (B) fraction of soluble AD brain extract after pre-exposure to monomer concentration ranging from 0 (100% binding), 0.08, 0.3, 1.25, and 5 μM. Percent binding response was calculated as: [(BRU) with monomers) / (BRU without monomers)] X100. Results shown are mean ± SEM of combined data from 6 independent studies for LMW and 3 independent studies for HMW. Not all antibodies were tested in all studies. (A) CREZ and GANT - 4 studies, PRX - 2 studies, LECAN, DONA, ADUC, ACU193 and PMN310 - 6 studies. (B) LECAN, DONA, ADUC, ACU193 and PMN310 - all 3 studies, CREZ, GANT and PRX - 1 study.

Based on the clinical experience to date, these results suggest a relationship between clinical benefit and the ability of Aβ-directed antibodies to avoid diversion by monomers and target toxic oligomers. The pan-reactive Aβ antibodies crenezumab and gantenerumab, which were highly susceptible to monomer competition, have failed in pivotal clinical trials [22, 23]. In contrast, antibodies such as aducanumab, lecanemab and donanemab, which retained binding to oligomers in the face of monomer competition, provided a clinical benefit [1–3]. Furthermore, plotting of the published clinical dementia rating scale sum of boxes (CDR-SB) results available from Phase 3 pivotal trials [1–3, 22, 23] showed a high degree of correlation between inhibition of cognitive decline and percent binding to LMW and HMW oligomers in the presence of monomer competition (R^2^ = 0.78 and 0.71, respectively) (Fig. 5).

**Figure 5.**
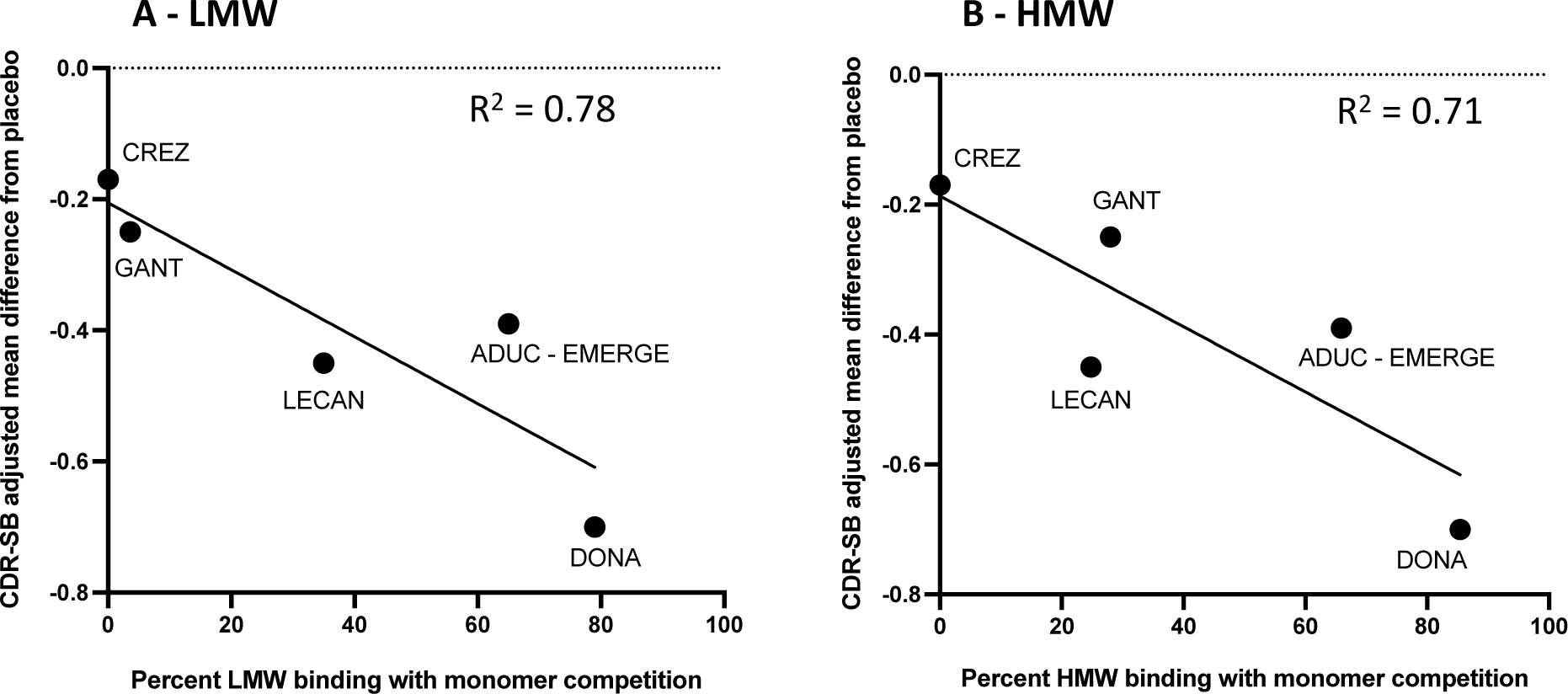
Correlation between CDR-SB benefit and percent binding to LMW and HMW oligomers in the presence of monomer competition. High degree of correlation observed between published CDR-SB adjusted mean difference from placebo (inhibition of cognitive decline) from pivotal Phase 3 clinical trials versus the mean percent binding of Aβ-directed antibodies to the LMW (A) or HMW (B) fraction of soluble AD brain extract after exposure to monomers (1.25 μM). Mean CDR-SB values of GRADUATE I and II trials for gantenerumab [22].

Along the same lines, a strong cognitive benefit was observed in preclinical studies with PMN310 which preferentially targets LMW oligomers and is highly resistant to monomer competition. Human APP [V717I] transgenic (hAPP-Tg) mice treated with a mouse IgG2a version of PMN310 starting at 5 months of age (30 mg/kg/week for 6 weeks) displayed significant protection of memory and learning compared to vehicle-treated hAPP-Tg mice in a Morris Water Maze test (Fig. 6). Both the escape latency (Fig. 6a) and search path (Fig. 6b) behaviors of PMN310-treated hAPP-Tg mice were comparable to those of non-Tg, wild-type mice.

**Figure 6.**
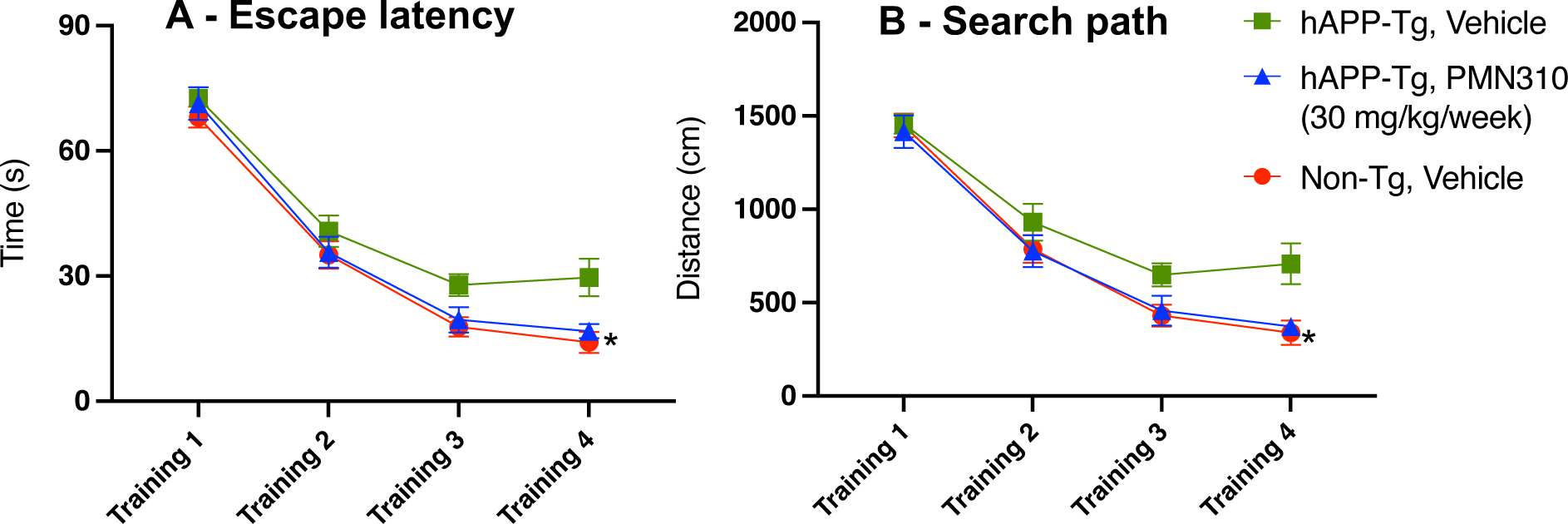
Cognitive benefit of PMN310 in AD mouse model. Human APP transgenic (hAPP-Tg) mice treated with a mouse IgG2a version of PMN310 showed significant protection of cognition as assessed in a Morris Water Maze test. (A) Escape latency was significantly affected by PMN310 treatment (*p*=0.0187; analyzed by two-way repeated measures ANOVA). Sidak’s post-hoc revealed a significant decrease in escape latency in the hAPP-Tg PMN310-treated mice compared to the hAPP-Tg vehicle-treated mice on training day 4 (*p*=0.0254). (B) Search path was also significantly affected by PMN310 treatment (*p*=0.0071; analyzed by two-way repeated measures ANOVA). Sidak’s post-hoc revealed a significant decrease in search path in the hAPP-Tg PMN310-treated mice compared to the hAPP-Tg vehicle-treated mice on training day 4 (*p*=0.0145).

### Binding of Aβ-directed antibodies to plaque

The reactivity of Aβ-directed antibodies with insoluble fibril deposits in AD brain sections was assessed by immunohistochemistry. Most antibodies (gantenerumab, aducanumab, lecanemab, donanemab, PRX h2731 and ACU193) showed staining of parenchymal plaque and vascular deposits of Aβ (Fig. 7). Where clinical data are available, binding to insoluble Aβ, in particular vascular deposits, has been associated with an increased incidence of amyloid-related imaging abnormalities (ARIA), in particular brain edema (ARIA-E), ranging from ∼13-35% depending on the antibody (Fig. 7) [1–3, 22, 23]. The underlying mechanism is believed to involve local inflammation triggered by the interaction of bound antibodies with complement [29] and Fc receptors on microglial cells leading to their activation [30]. The consequent release of pro-inflammatory mediators, together with potential weakening of the vascular wall following amyloid removal, is thought to result in ARIA-E and microhemorrhages (ARIA-H) [29, 31–34]. In agreement with this premise, solanezumab, which targets monomers and does not bind plaque (Fig. 7), did not produce an increased incidence of ARIA [35]. Notably, PMN310, which is selective for soluble toxic Aβ oligomers, did not produce any detectable staining of plaque or vascular deposits in AD brain sections (Fig. 7). The ability of PMN310 to avoid both monomer and plaque binding would be expected to increase the available effective dose against toxic oligomers. Importantly, it may also reduce the risk of ARIA-E and ARIA-H observed with other Aβ-directed antibodies. This premise is supported by preclinical toxicology studies in which weekly dosing of a murine IgG2a version of PMN310 in plaque-bearing knock-in APP^SAA^ mice [36] at 800 mg/kg for 26 weeks did not cause brain hemorrhages upon microscopic examination using Perls’ Prussian Blue staining to detect the presence of hemosiderin (Fig. 8).

**Figure 7.**
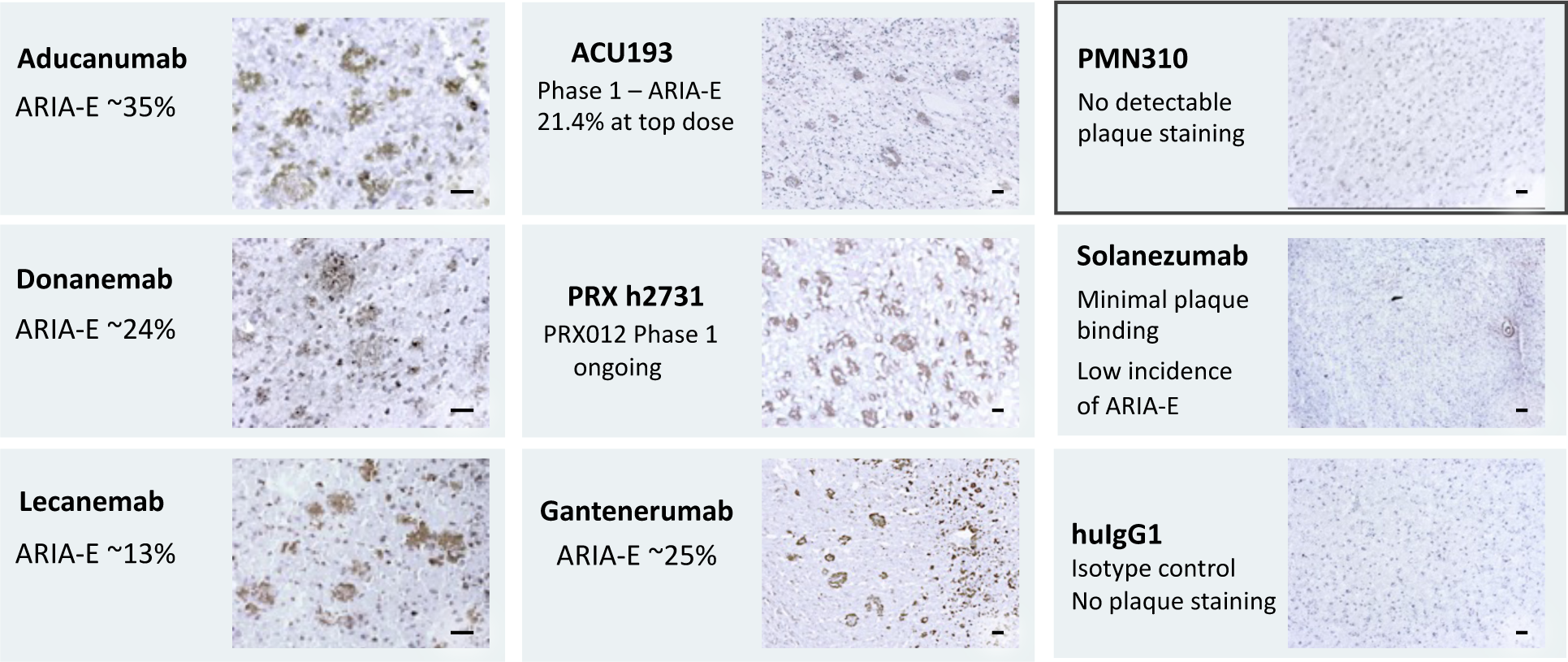
Reactivity of Aβ-directed antibodies with insoluble fibril deposits in AD brain. Representative images of plaque staining in brain sections from AD frontal cortex. All antibodies were tested at the same concentration of 1 μg/ml. Scale bars = 50 μm.

**Figure 8.**
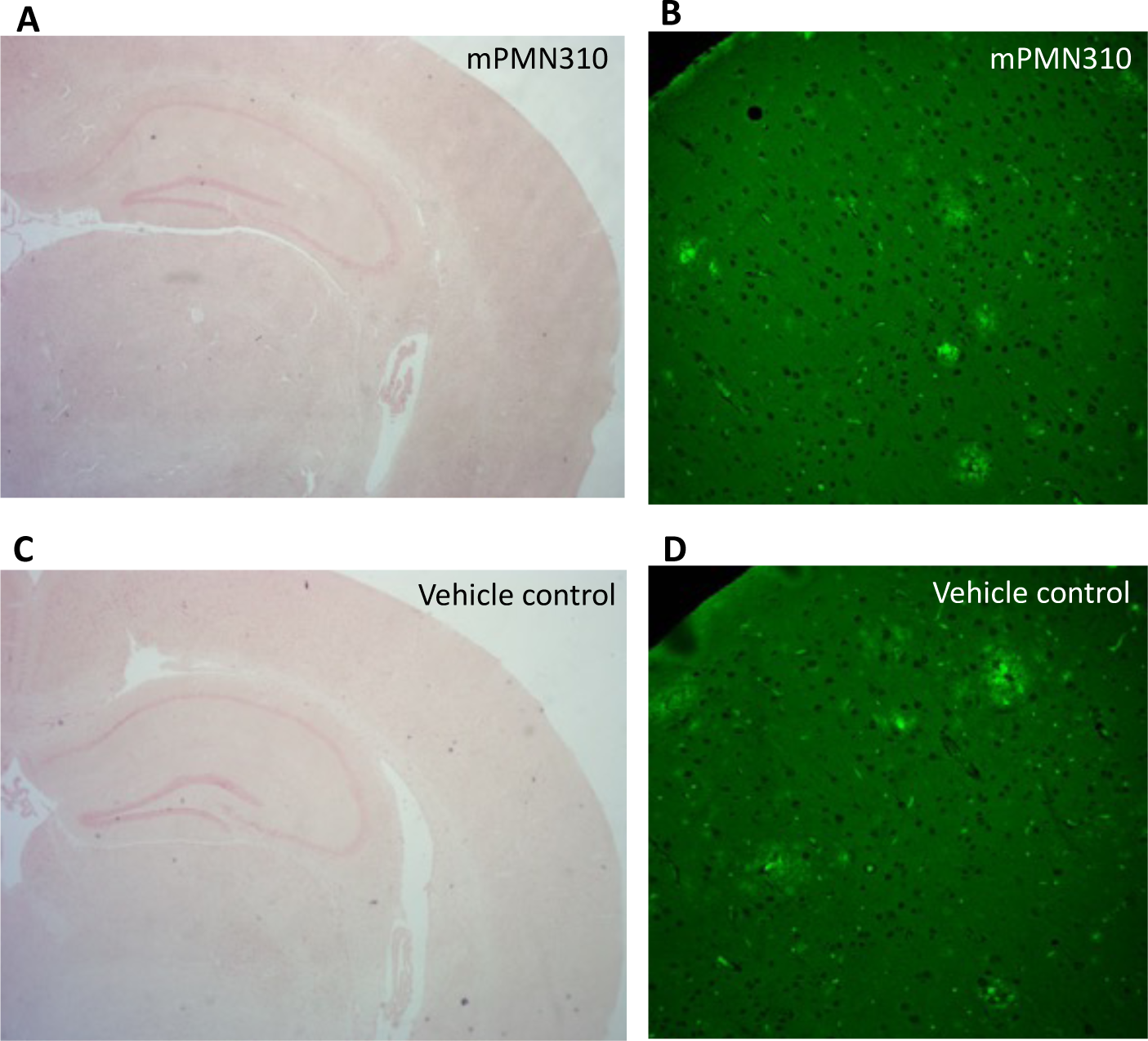
Treatment of *APP^SAA^*mice at 800 mg/kg/week for 26 weeks does not cause brain hemorrhages. Brain sections from mice treated with murine PMN310 (A, B) or vehicle (C, D) were stained with Perls’ Prussian Blue stain to detect the presence of hemosiderin as an indicator of hemorrhage in the brain (A, C). Stained sections from 29 vehicle control mice and 29 murine PMN310-treated mice were examined (14 males and 15 females in each group). The presence of plaque was confirmed by Amylo-Glo staining (B, D). Representative images at 25X magnification are shown. An unrelated stock spleen sample was used as a positive control to validate the Perls’ Prussian Blue staining procedure (Supplemental Figure 1).

## DISCUSSION

As progress continues in the development of therapeutic Aβ-directed antibodies for AD, it has become increasingly clear that efficacy is dependent on the species of Aβ being targeted. Targeting monomers with solanezumab failed to impact cognitive decline [37] and it has been suggested that maintaining high levels of Aβ42 monomers in the CNS may actually benefit synaptic plasticity and memory [38]. This contention is supported by an observed positive correlation between high levels of Aβ42 in the CSF and normal cognition in amyloid-positive individuals with AD-causing genetic mutations [38]. In a related finding, treatment of AD patients with BACE-1 inhibitors, which dramatically reduced levels of monomers in the CSF, resulted in worsening of cognition [39]. Antibody-mediated plaque clearance has been more successful but has produced only a modest inhibition of cognitive decline (∼20-30%, [1–3]) accompanied by an increased risk of ARIA. A large body of experimental and clinical evidence currently points to soluble toxic Aβ aggregates, rather than insoluble fibrils and plaque, as the primary driver of disease in AD patients [5–10].

As described here, toxic soluble Aβ aggregates likely exist in the brain as a continuum of oligomers and protofibrils of various sizes and shapes as opposed to discrete populations with defined molecular weights. Based on the observed relationship between clinical efficacy and the ability to bind oligomers in the face of monomer competition, we propose that the efficacy of Aβ-directed antibodies may depend on achieving an effective dose capable of clearing soluble toxic species spanning a range of molecular weights despite the target distraction of soluble monomers. This premise is consistent with the clinical experience to date showing a strong correlation between CDR-SB benefit (inhibition of cognitive decline) in pivotal Phase 3 clinical trials (Fig. 5) and the percent binding of Aβ-directed antibodies to LMW and HMW oligomers in the presence of monomer competition measured in this study (Fig. 4). A correlation between plaque clearance and CDR-SB benefit has also been reported [4]. However, all plaque binding antibodies in this study also bind oligomers, and, at the same time, clinical data indicate that plaque clearance does not appear to be sufficient to inhibit disease progression. For example, gantenerumab failed to achieve a significant clinical benefit despite its ability to remove plaque [22]. Similarly, treatment gap data collected between trial completion and initiation of an open label extension (OLE) with lecanemab and aducanumab have shown worsening of cognition after cessation of treatment even though plaque burden had been reduced and remained flat during the gap period [40, 41]. These results suggest that removal of soluble toxic oligomers may be the primary driver of clinical benefit and must be sustained for continued efficacy.

In earlier stages of clinical development, Phase 1 clinical testing of ACU193, an antibody reported to have approximately 500-fold greater selectivity for Aβ oligomers over monomers and more than 85-fold greater selectivity for oligomers *vs* fibrillar Aβ, showed encouraging amelioration of biomarkers associated with AD pathology (pTau181, neurogranin, Aβ42/40 ratio, VAMP2, GFAP) after 3 months of dosing in patients with early AD [42]. Plaque reduction was observed in the high-dose cohorts (60 mg/kg Q4W and 25 mg/kg Q2W) likely due to direct binding of ACU193 to plaque (Fig. 7) or potentially involving other mechanisms such as a shift in the oligomer-fibril equilibrium or a change in the milieu favoring microglial removal of plaque upon neutralization of inflammatory oligomers. Treatment with ACU193 was associated with a 21.4% incidence of ARIA-E at the highest dose of 60 mg/kg (combined single ascending dose and multiple ascending dose groups), a much higher dose than that used with other plaque-binding antibodies. The use of a human IgG2 isotype with reduced effector function *vs* the human IgG1 effector isotype of other antibodies may potentially have played a role in dampening inflammatory responses that can trigger ARIA following binding to Aβ plaque/vascular deposits. Phase 1 testing of the PRX012 antibody is ongoing and results are pending at this time. The impact of ACU193 and PRX012 treatment on cognition and relationship with their binding profile remain to be determined.

Of all the antibodies tested in this study, only PMN310 selectively targeted oligomers without cross-reacting with monomers or plaque (Figs. 2, 3, 7, Table 1). The ability of PMN310 to avoid both monomer and plaque binding may result in greater potency by increasing the proportion of dosed antibody available to target toxic oligomers. In preclinical mouse studies, targeting of AβO by PMN310 resulted in the protection of cognition in hAPP Tg mice (Fig. 6) and, as previously reported, in mice injected intracerebroventricularly (ICV) with AβO [14]. Importantly, based on the current understanding of the biology of ARIA implicating binding of antibodies to Aβ deposits as an initiating factor, PMN310 may also present a reduced risk of ARIA-E and ARIA-H compared to other Aβ-directed antibodies. In agreement with this concept, intensive treatment of plaque-bearing APP^SAA^ mice with high doses of a mouse version of PMN310 (800 mg/kg/week for 26 weeks) did not produce microhemorrhages (Fig. 8). In contrast, treatment of transgenic AD mice with plaque-binding antibodies including bapineuzumab, gantenerumab and aducanumab has been shown to induce microhemorrhages with less stringent dosing regimens [30, 31, 34, 43, 44]. Clinical Phase 1a testing of PMN310 in normal healthy volunteers was initiated in November 2023 (NCT06105528). Subsequent testing in AD patients may provide the basis for a true test of the “oligomer hypothesis” without confounding cross-reactivity with other Aβ species.

**Table 1.**
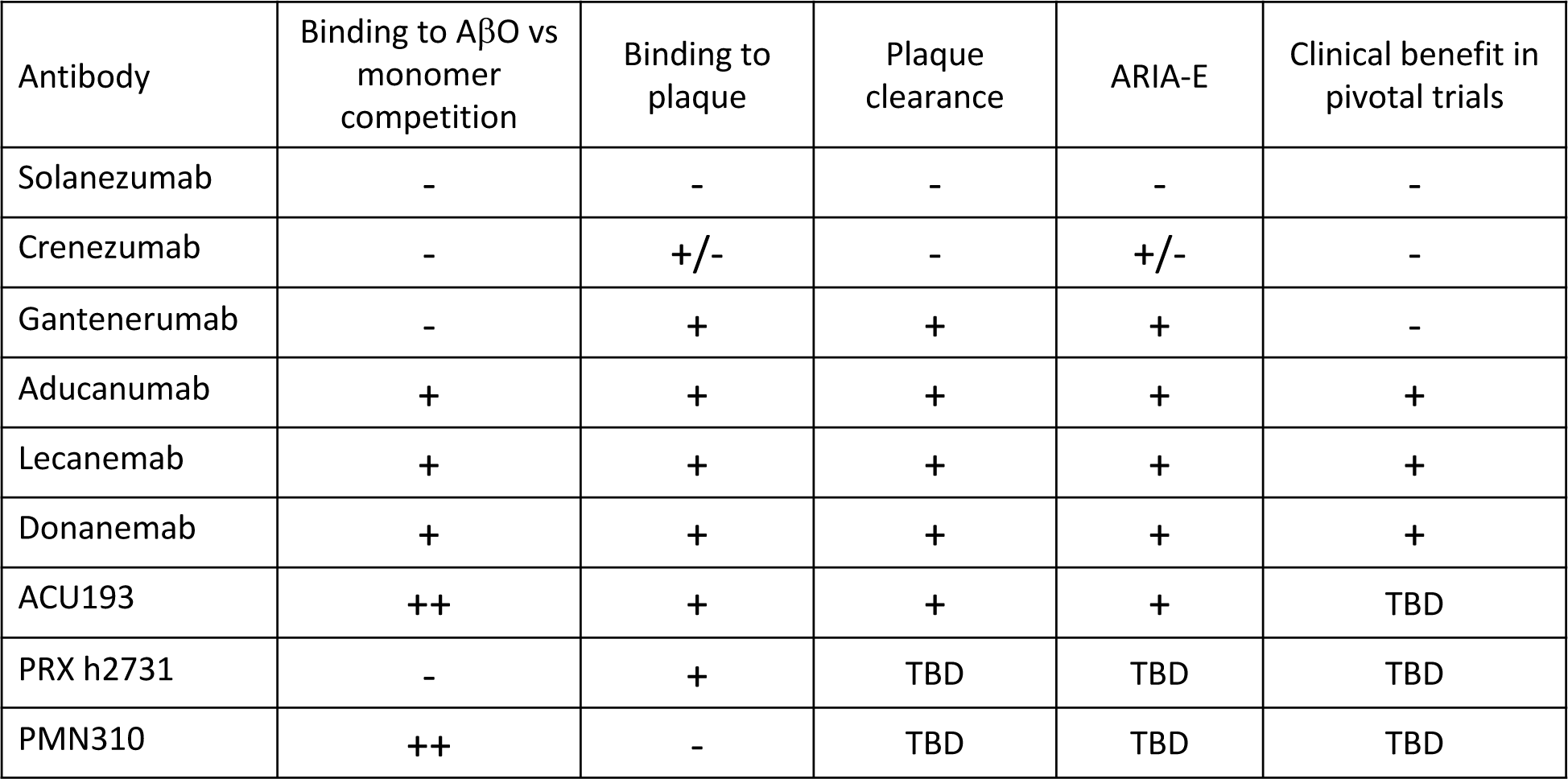
Summary of findings and clinical experience. TBD: To be determined

## Author contributions

Johanne M Kaplan (conceptualization, data curation, formal analysis, project administration, writing – original draft & editing); Ebrima Gibbs (conceptualization, formal analysis, investigation, methodology, review & editing); Juliane Coutts (investigation, methodology, visualization, review & editing), Beibei Zhao (conceptualization, review & editing), Ian Mackenzie (resources, review & editing), Neil R Cashman (conceptualization, supervision, funding acquisition, review & editing)

## Acknowledgments

The authors wish to thank Dragana Vidakovic for technical support and Dr. Larry Altsteil for critical review of the manuscript.

## Data availability statement

The data supporting the findings of this study are available within the article.

---

**Supplemental Figure 1.**
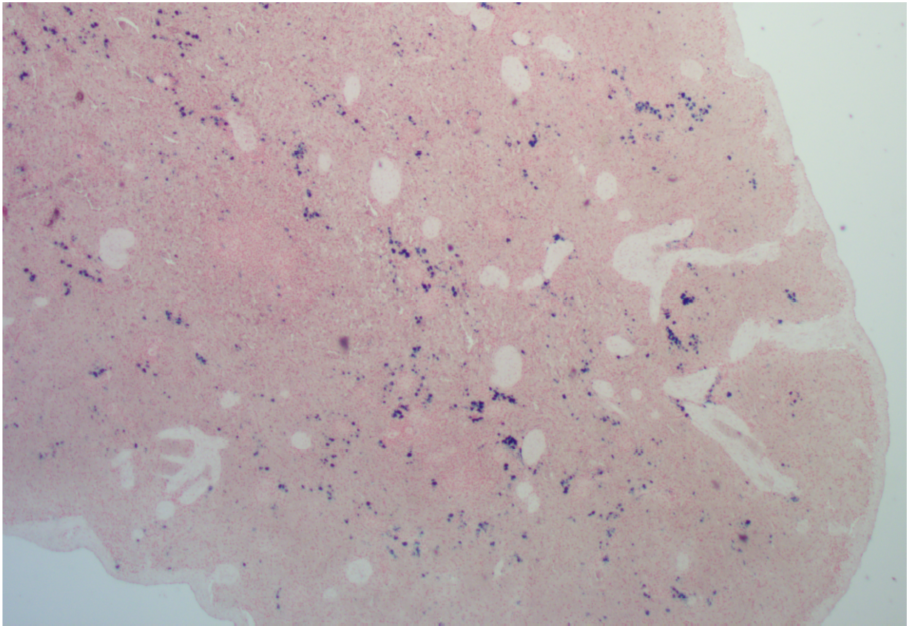
Perls’ Prussian Blue staining of a stock spleen sample confirmed the validity of the staining procedure.

**Supplemental Table 1.** AD brain samples. PMI: Post-mortem interval, N/A: Not available

| Sample ID number | Sex | Age | PMI |
| --- | --- | --- | --- |
| NDBB230 | M | 74 | 12-36 h |
| NDBB232 | F | 75 | 12-36 h |
| NIH 5682 | M | 68 | 8 h |
| NDBB029 | M | 73 | < 24 h |
| NDBB084 | M | 77 | < 24 h |
| NDBB093 | F | 81 | < 48 h |
| NDBB167 | F | 76 | < 36 h |
| NDBB170 | M | 62 | N/A |
| NDBB054 | M | 83 | 24 h |
| NDBB097 | M | 70 | 24-48 h |
| NDBB121 | F | 55 | < 24 h |
| NDBB185 | M | 59 | N/A |
| NDBB024 | M | 73 | < 24 h |
| NDBB059 | F | 60 | 48 h |
| NIH5598 | M | 80 | 5 h |

PMI: Post-mortem interval, N/A: Not available

## REFERENCES

[1] Budd Haeberlein S, Aisen PS, Barkhof F, Chalkias S, Chen T, Cohen S, Dent G, Hansson O, Harrison K, von Hehn C, Iwatsubo T, Mallinckrodt C, Mummery CJ, Muralidharan KK, Nestorov I, Nisenbaum L, Rajagovindan R, Skordos L, Tian Y, van Dyck CH, Vellas B, Wu S, Zhu Y, Sandrock A (2022) Two Randomized Phase 3 Studies of Aducanumab in Early Alzheimer’s Disease. J Prev Alzheimers Dis 9, 197–210.

[2] Sims JR, Zimmer JA, Evans CD, Lu M, Ardayfio P, Sparks J, Wessels AM, Shcherbinin S, Wang H, Monkul Nery ES, Collins EC, Solomon P, Salloway S, Apostolova LG, Hansson O, Ritchie C, Brooks DA, Mintun M, Skovronsky DM, Investigators T-A (2023) Donanemab in Early Symptomatic Alzheimer Disease: The TRAILBLAZER-ALZ 2 Randomized Clinical Trial. JAMA 330, 512–527.

[3] van Dyck CH, Swanson CJ, Aisen P, Bateman RJ, Chen C, Gee M, Kanekiyo M, Li D, Reyderman L, Cohen S, Froelich L, Katayama S, Sabbagh M, Vellas B, Watson D, Dhadda S, Irizarry M, Kramer LD, Iwatsubo T (2023) Lecanemab in Early Alzheimer’s Disease. N Engl J Med 388, 9–21.

[4] Boxer AL, Sperling R (2023) Accelerating Alzheimer’s therapeutic development: The past and future of clinical trials. Cell 186, 4757–4772.

[5] Balducci C, Beeg M, Stravalaci M, Bastone A, Sclip A, Biasini E, Tapella L, Colombo L, Manzoni C, Borsello T, Chiesa R, Gobbi M, Salmona M, Forloni G (2010) Synthetic amyloid-beta oligomers impair long-term memory independently of cellular prion protein. Proc Natl Acad Sci U S A 107, 2295–2300.

[6] Cleary JP, Walsh DM, Hofmeister JJ, Shankar GM, Kuskowski MA, Selkoe DJ, Ashe KH (2005) Natural oligomers of the amyloid-beta protein specifically disrupt cognitive function. Nat Neurosci 8, 79–84.

[7] Cline EN, Bicca MA, Viola KL, Klein WL (2018) The Amyloid-beta Oligomer Hypothesis: Beginning of the Third Decade. J Alzheimers Dis 64, S567–S610.

[8] Ferreira ST, Lourenco MV, Oliveira MM, De Felice FG (2015) Soluble amyloid-beta oligomers as synaptotoxins leading to cognitive impairment in Alzheimer’s disease. Front Cell Neurosci 9, 191.

[9] Shankar GM, Li S, Mehta TH, Garcia-Munoz A, Shepardson NE, Smith I, Brett FM, Farrell MA, Rowan MJ, Lemere CA, Regan CM, Walsh DM, Sabatini BL, Selkoe DJ (2008) Amyloid-beta protein dimers isolated directly from Alzheimer’s brains impair synaptic plasticity and memory. Nat Med 14, 837–842.

[10] Stohr J, Watts JC, Mensinger ZL, Oehler A, Grillo SK, DeArmond SJ, Prusiner SB, Giles K (2012) Purified and synthetic Alzheimer’s amyloid beta (Abeta) prions. Proc Natl Acad Sci U S A 109, 11025–11030.

[11] Sun H, Wang Y, Wang Y, Zeng F (2024) Case of Autosomal Dominant Alzheimer Disease With Negative Findings From PiB-PET Examination. Neurol Genet 10, e200119.

[12] Tomiyama T, Shimada H (2020) APP Osaka Mutation in Familial Alzheimer’s Disease-Its Discovery, Phenotypes, and Mechanism of Recessive Inheritance. Int J Mol Sci 21.

[13] Scholl M, Wall A, Thordardottir S, Ferreira D, Bogdanovic N, Langstrom B, Almkvist O, Graff C, Nordberg A (2012) Low PiB PET retention in presence of pathologic CSF biomarkers in Arctic APP mutation carriers. Neurology 79, 229–236.

[14] Gibbs E, Silverman JM, Zhao B, Peng X, Wang J, Wellington CL, Mackenzie IR, Plotkin SS, Kaplan JM, Cashman NR (2019) A Rationally Designed Humanized Antibody Selective for Amyloid Beta Oligomers in Alzheimer’s Disease. Sci Rep 9, 9870.

[15] Benilova I, Karran E, De Strooper B (2012) The toxic Abeta oligomer and Alzheimer’s disease: an emperor in need of clothes. Nat Neurosci 15, 349–357.

[16] Goure WF, Krak GA, Jerecic J, Hefti F (2014) Targeting the proper amyloid-beta neuronal toxins: a path forward for Alzheimer’s disease immunotherapeutics. Alzheimers Res Ther 6, 42.

[17] Tolar M, Hey J, Power A, Abushakra S (2021) Neurotoxic Soluble Amyloid Oligomers Drive Alzheimer’s Pathogenesis and Represent a Clinically Validated Target for Slowing Disease Progression. Int J Mol Sci 22.

[18] Gandy S, Ehrlich ME (2023) Moving the Needle on Alzheimer’s Disease with an Anti-Oligomer Antibody. N Engl J Med 388, 80–81.

[19] Lannfelt L, Söderberg, L., Laudon, H., Sahlin, C., Johannesson, M., Nygren, P., Möller, C. (2019) BAN2401 shows stronger binding to soluble aggregated amyloid-beta species than aducanumab. In AAIC, Philadelphia, USA. https://bioarctic.se/sv/wp-content/uploads/sites/4/2019/07/ban2401-poster-aaic-july-2019-los-angeles-usa.pdf

[20] Sehlin D, Englund H, Simu B, Karlsson M, Ingelsson M, Nikolajeff F, Lannfelt L, Pettersson FE (2012) Large aggregates are the major soluble Abeta species in AD brain fractionated with density gradient ultracentrifugation. PLoS One 7, e32014.

[21] Soderberg L, Johannesson M, Nygren P, Laudon H, Eriksson F, Osswald G, Moller C, Lannfelt L (2023) Lecanemab, Aducanumab, and Gantenerumab - Binding Profiles to Different Forms of Amyloid-Beta Might Explain Efficacy and Side Effects in Clinical Trials for Alzheimer’s Disease. Neurotherapeutics 20, 195–206.

[22] Bateman RJ, Smith J, Donohue MC, Delmar P, Abbas R, Salloway S, Wojtowicz J, Blennow K, Bittner T, Black SE, Klein G, Boada M, Grimmer T, Tamaoka A, Perry RJ, Turner RS, Watson D, Woodward M, Thanasopoulou A, Lane C, Baudler M, Fox NC, Cummings JL, Fontoura P, Doody RS, Graduate I, Investigators II, the Gantenerumab Study G (2023) Two Phase 3 Trials of Gantenerumab in Early Alzheimer’s Disease. N Engl J Med 389, 1862–1876.

[23] Ostrowitzki S, Bittner T, Sink KM, Mackey H, Rabe C, Honig LS, Cassetta E, Woodward M, Boada M, van Dyck CH, Grimmer T, Selkoe DJ, Schneider A, Blondeau K, Hu N, Quartino A, Clayton D, Dolton M, Dang Y, Ostaszewski B, Sanabria-Bohorquez SM, Rabbia M, Toth B, Eichenlaub U, Smith J, Honigberg LA, Doody RS (2022) Evaluating the Safety and Efficacy of Crenezumab vs Placebo in Adults With Early Alzheimer Disease: Two Phase 3 Randomized Placebo-Controlled Trials. JAMA Neurol 79, 1113–1121.

[24] Holtta M, Hansson O, Andreasson U, Hertze J, Minthon L, Nagga K, Andreasen N, Zetterberg H, Blennow K (2013) Evaluating amyloid-beta oligomers in cerebrospinal fluid as a biomarker for Alzheimer’s disease. PLoS One 8, e66381.

[25] Kass B, Schemmert S, Zafiu C, Pils M, Bannach O, Kutzsche J, Bujnicki T, Willbold D (2022) Abeta oligomer concentration in mouse and human brain and its drug-induced reduction ex vivo. Cell Rep Med 3, 100630.

[26] Savage MJ, Kalinina J, Wolfe A, Tugusheva K, Korn R, Cash-Mason T, Maxwell JW, Hatcher NG, Haugabook SJ, Wu G, Howell BJ, Renger JJ, Shughrue PJ, McCampbell A (2014) A sensitive abeta oligomer assay discriminates Alzheimer’s and aged control cerebrospinal fluid. J Neurosci 34, 2884–2897.

[27] Reed MN, Hofmeister JJ, Jungbauer L, Welzel AT, Yu C, Sherman MA, Lesne S, LaDu MJ, Walsh DM, Ashe KH, Cleary JP (2011) Cognitive effects of cell-derived and synthetically derived Abeta oligomers. Neurobiol Aging 32, 1784–1794.

[28] Yang T, Li S, Xu H, Walsh DM, Selkoe DJ (2017) Large Soluble Oligomers of Amyloid beta-Protein from Alzheimer Brain Are Far Less Neuroactive Than the Smaller Oligomers to Which They Dissociate. J Neurosci 37, 152–163.

[29] Lemere CA (2023) Is ARIA an Inflammatory Reaction to Vascular Amyloid? In AAIC, Amsterdam, Netherlands. https://www.alzforum.org/news/conference-coverage/aria-inflammatory-reaction-vascular-amyloid

[30] Bussière T (2024) Mouse Models and Markers for Cerebral Amyloid Angiopathy, ARIA. In AD/PD 2024. https://www.alzforum.org/news/conference-coverage/mouse-models-and-markers-cerebral-amyloid-angiopathy-aria

[31] Pfeifer M, Boncristiano S, Bondolfi L, Stalder A, Deller T, Staufenbiel M, Mathews PM, Jucker M (2002) Cerebral hemorrhage after passive anti-Abeta immunotherapy. Science 298, 1379.

[32] Roytman M, Mashriqi F, Al-Tawil K, Schulz PE, Zaharchuk G, Benzinger TLS, Franceschi AM (2023) Amyloid-Related Imaging Abnormalities: An Update. AJR Am J Roentgenol 220, 562–574.

[33] Sperling RA, Aisen PS, Beckett LA, Bennett DA, Craft S, Fagan AM, Iwatsubo T, Jack CR, Jr., Kaye J, Montine TJ, Park DC, Reiman EM, Rowe CC, Siemers E, Stern Y, Yaffe K, Carrillo MC, Thies B, Morrison-Bogorad M, Wagster MV, Phelps CH (2011) Toward defining the preclinical stages of Alzheimer’s disease: recommendations from the National Institute on Aging-Alzheimer’s Association workgroups on diagnostic guidelines for Alzheimer’s disease. Alzheimers Dement 7, 280–292.

[34] Taylor X, Clark IM, Fitzgerald GJ, Oluoch H, Hole JT, DeMattos RB, Wang Y, Pan F (2023) Amyloid-beta (Abeta) immunotherapy induced microhemorrhages are associated with activated perivascular macrophages and peripheral monocyte recruitment in Alzheimer’s disease mice. Mol Neurodegener 18, 59.

[35] Carlson C, Siemers E, Hake A, Case M, Hayduk R, Suhy J, Oh J, Barakos J (2016) Amyloid-related imaging abnormalities from trials of solanezumab for Alzheimer’s disease. Alzheimers Dement (Amst) 2, 75–85.

[36] Xia D, Lianoglou S, Sandmann T, Calvert M, Suh JH, Thomsen E, Dugas J, Pizzo ME, DeVos SL, Earr TK, Lin CC, Davis S, Ha C, Leung AW, Nguyen H, Chau R, Yulyaningsih E, Lopez I, Solanoy H, Masoud ST, Liang CC, Lin K, Astarita G, Khoury N, Zuchero JY, Thorne RG, Shen K, Miller S, Palop JJ, Garceau D, Sasner M, Whitesell JD, Harris JA, Hummel S, Gnorich J, Wind K, Kunze L, Zatcepin A, Brendel M, Willem M, Haass C, Barnett D, Zimmer TS, Orr AG, Scearce-Levie K, Lewcock JW, Di Paolo G, Sanchez PE (2022) Novel App knock-in mouse model shows key features of amyloid pathology and reveals profound metabolic dysregulation of microglia. Mol Neurodegener 17, 41.

[37] Honig LS, Vellas B, Woodward M, Boada M, Bullock R, Borrie M, Hager K, Andreasen N, Scarpini E, Liu-Seifert H, Case M, Dean RA, Hake A, Sundell K, Poole Hoffmann V, Carlson C, Khanna R, Mintun M, DeMattos R, Selzler KJ, Siemers E (2018) Trial of Solanezumab for Mild Dementia Due to Alzheimer’s Disease. N Engl J Med 378, 321–330.

[38] Sturchio A, Dwivedi AK, Malm T, Wood MJA, Cilia R, Sharma JS, Hill EJ, Schneider LS, Graff-Radford NR, Mori H, Nubling G, El Andaloussi S, Svenningsson P, Ezzat K, Espay AJ, Dominantly Inherited Alzheimer C (2022) High Soluble Amyloid-beta42 Predicts Normal Cognition in Amyloid-Positive Individuals with Alzheimer’s Disease-Causing Mutations. J Alzheimers Dis 90, 333–348.

[39] Egan MF, Kost J, Voss T, Mukai Y, Aisen PS, Cummings JL, Tariot PN, Vellas B, van Dyck CH, Boada M, Zhang Y, Li W, Furtek C, Mahoney E, Harper Mozley L, Mo Y, Sur C, Michelson D (2019) Randomized Trial of Verubecestat for Prodromal Alzheimer’s Disease. N Engl J Med 380, 1408–1420.

[40] Cohen S, van Dyck, C. H., Mummery, C. J., Porsteinsson, A., Kong, J., Miller, R., Racine, A., O’Gorman, J., Haeberlein, S. B., Salloway, S. (2021) Baseline EMBARK data from EMERGE, ENGAGE and PRIME participants in the EMBARK re-dosing study. In CTAD, Boston, USA. https://investors.biogen.com/static-files/2fe89f45-2ed2-4d5d-92ca-ac5003058ce2

[41] McDade E, Cummings JL, Dhadda S, Swanson CJ, Reyderman L, Kanekiyo M, Koyama A, Irizarry M, Kramer LD, Bateman RJ (2022) Lecanemab in patients with early Alzheimer’s disease: detailed results on biomarker, cognitive, and clinical effects from the randomized and open-label extension of the phase 2 proof-of-concept study. Alzheimers Res Ther 14, 191.

[42] Trame M, Siemers, E., Salloway, S., Trame, M. N. (2023) INTERCEPT-AD phase 1 insights and findings from the investigation of ACU193, a monoclonal antibody targeting soluble Aβ oligomers. In CTAD, Boston, USA. https://investors.acumenpharm.com/static-files/e9f30ca7-82e1-463d-bc94-8932c76f4b41

[43] Luo F, Rustay NR, Seifert T, Roesner B, Hradil V, Hillen H, Ebert U, Severin JM, Cox BF, Llano DA, Day M, Fox GB (2010) Magnetic resonance imaging detection and time course of cerebral microhemorrhages during passive immunotherapy in living amyloid precursor protein transgenic mice. J Pharmacol Exp Ther 335, 580–588.

[44] Racke MM, Boone LI, Hepburn DL, Parsadainian M, Bryan MT, Ness DK, Piroozi KS, Jordan WH, Brown DD, Hoffman WP, Holtzman DM, Bales KR, Gitter BD, May PC, Paul SM, DeMattos RB (2005) Exacerbation of cerebral amyloid angiopathy-associated microhemorrhage in amyloid precursor protein transgenic mice by immunotherapy is dependent on antibody recognition of deposited forms of amyloid beta. J Neurosci 25, 629–636.

